# Corticostriatal ensemble dynamics across heroin self-administration to reinstatement

**DOI:** 10.1101/2024.06.21.599790

**Authors:** Rachel E. Clarke, Roger I. Grant, Shannon N. Woods, Bayleigh E. Pagoota, Sophie Buchmaier, Bogdan Bordieanu, Anna Tsyrulnikov, Annaka M. Westphal, Jacqueline E Paniccia, Elizabeth M Doncheck, Jayda Carroll-Deaton, Kelsey M Vollmer, Amy L. Ward, Kion T. Winston, Danielle I. King, Jade Baek, Mike R. Martino, Lisa M. Green, Jacqueline F. McGinty, Michael D. Scofield, James M. Otis

## Abstract

Corticostriatal projection neurons from prelimbic medial prefrontal cortex to the nucleus accumbens core critically regulate drug-seeking behaviors, yet the underlying encoding dynamics whereby these neurons contribute to drug seeking remain elusive. Here we use two-photon calcium imaging to visualize the activity of corticostriatal neurons in mice from the onset of heroin use to relapse. We find that the activity of these neurons is highly heterogeneous during heroin self-administration and seeking, with at least 8 distinct neuronal ensembles that display both excitatory and inhibitory encoding dynamics. These neuronal ensembles are particularly apparent during relapse, where excitatory responses are amplified compared to heroin self-administration. Moreover, we find that optogenetic inhibition of corticostriatal projection neurons attenuates heroin seeking regardless of the relapse trigger. Our results reveal the precise corticostriatal activity dynamics underlying drug-seeking behaviors and support a key role for this circuit in mediating relapse to drug seeking.

## INTRODUCTION

Substance use disorder is a chronically relapsing disorder, characterized by long lasting neurobiological adaptations in brain regions that encode reward^1^. One hallmark of addiction is dysregulated prefrontal cortex activity that manifests as hypoactivity at baseline, and hyperactivity in response to drug-associated cues^2,3^. Despite clear evidence that substance abuse results in aberrant prefrontal cortex activity, leading to impaired behavioral inhibition of maladaptive drug-seeking^2^, we currently lack an effective treatment strategy to target the neurobiological adaptations to substance abuse within the prefrontal cortex and prevent relapse in the long-term.

The extreme heterogeneity of activity dynamics, gene expression, projection targets, and afferent connectivity that exists within the prefrontal cortex means it is a challenging brain region to study and target. For example, there is now converging evidence from human and preclinical studies that supports a critical role for the corticostriatal circuit connecting the prelimbic medial prefrontal cortex (PrL) projections to the nucleus accumbens core (NAcc) in drug seeking^4–14^. However, even within circuit-specific projection neurons there is heterogeneity of activity dynamics^15^ and cell types^16,17^ making it difficult to decipher the necessary components of prefrontal cortex activity that are required to initiate and maintain drug seeking. To date, no studies have investigated the precise activity dynamics of the neuronal ensembles within the corticostriatal projection neurons during drug seeking, representing a vital gap in our understanding of how activity in this circuit contributes to drug seeking. To address this issue, here we use a newly developed head-fixed model of heroin self-administration in mice^18,19^, enabling simultaneous two-photon calcium imaging of PrL→NAcc neurons from the onset of heroin self-administration through reinstatement (a model of relapse).

## RESULTS

### Optogenetic inhibition of PrL→NAcc circuit prevents cue-, drug- and stress-induced heroin seeking

Previous studies have demonstrated glutamatergic PrL→NAcc projections are necessary for cue- and drug-induced reinstatement of heroin seeking^7,8^. However, these studies were performed in freely moving rodents, and whether this circuit is required for heroin seeking in a head-fixed model of heroin self-administration has not been tested. Moreover, whether PrL→NAcc projection neurons are required for stress-induced heroin seeking is unknown. Therefore, we used optogenetics to inhibit PrL→NAcc neuronal activity during cue-, drug- and stress-induced heroin seeking using our previously reported model of head-fixed heroin self-administration^18,19^. Firstly, we trained head-fixed mice (Fig. 1A) to press an active, but not inactive, lever for presentation of an auditory cue preceding subsequent delivery of heroin (Fig. 1B). Total active lever presses were used as an index of drug seeking, and goal-directed behavior assessed by lever discrimination (active vs inactive lever presses). Mice underwent 14 days of heroin self-administration (Fig. 1C) followed immediately by 10 days of extinction training, wherein active lever presses no longer resulted in heroin or cue delivery resulting in an attenuation of active lever pressing compared to early extinction (Fig. 1D). To acutely inhibit PrL→NAcc projection neurons, we delivered a retrogradely-trafficked virus encoding Cre-recombinase (rgAAV2-CAG-Cre) bilaterally into the NAcc and a Cre-dependent virus encoding halorhodopsin (AAV5-DIO-Ef1α-eNpHR3.0-eYFP) or control enhanced yellow fluorescent protein (AAV5-ef1α-DIO-eYFP) into PrL, with optical fibers implanted dorsal to PrL (Fig. 1E-F, Fig. S1A). To model “relapse” to drug seeking, after self-administration and extinction mice underwent reinstatement testing where they were re-exposed to the heroin-associated auditory cue (cue-reinstatement), given a priming injection of heroin (1 mg/kg, ip; drug-reinstatement), or exposed to predator odor (15 min pre-session 2,5-dihydro-2,4,5-trimethylthiazoline, TMT; stress-reinstatement). Optogenetic inhibition of the PrL→NAcc projection neurons during cue, drug- and stress-induced reinstatement tests abolished active lever pressing (Fig. 1G), with no differences observed between halorhodopsin and eYFP groups under laser off conditions (Fig S1B), or in inactive lever pressing rates (Fig S1C-D). Therefore, our results suggest that PrL→NAcc projection neurons are necessary for cue-, drug- and stress-induced reinstatement of heroin seeking. However, the precise activity dynamics of the PrL→NAcc projection neurons required for reinstatement of heroin seeking remain unknown.

**Figure 1.**
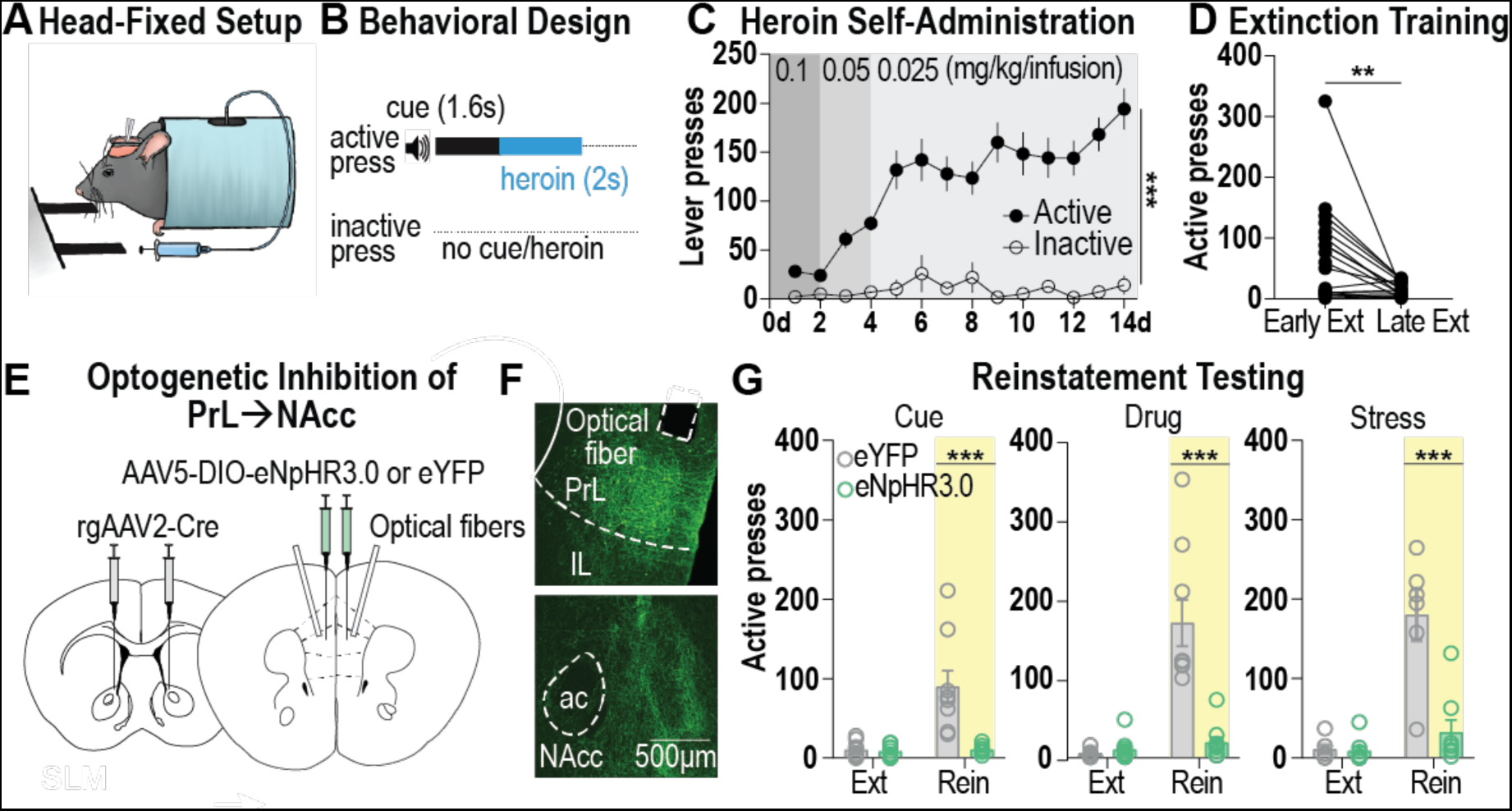
PrL→NAcc circuit is necessary for reinstatement of heroin seeking. (**A**) Schematic showing head-fixed behavioral apparatus. (**B**) Behavioral paradigm for intravenous head-fixed heroin self-administration. (**C**) Grouped data for acquisition of heroin self-administration. Mice learn to discriminate between the active and inactive levers, with greater active lever presses across acquisition (n=19 mice; lever: F_1,36_ = 119.7, p<0.001). (**D**) Grouped data for extinction training. Mice decreased activer lever pressing following a minimum of 10 days of extinction training (n =19 mice; averaged first 3-days of extinction vs last 3 days of extinction: t_18_=3.39, p=0.003). (**E**) Viral approach used to optogenetically inhibit PrL neurons that project to the NAcc. (**F**) Representative images showing eNpHR3.0-eYFP expression and optical fiber placement in the PrL (top), and eNpHR3.0-eYFP expressing fibers in the NAcc (bottom). (**G**) Optogenetic inhibition of PrL→NAcc neurons prevents cue-, drug- and stress-induced reinstatement of heroin seeking, measured as active lever presses (n= 6-10/group; cue reinstatement: interaction: F_1,17_=21.36, p<0.001, group comparisons: ext: p=0.9935, rein: p<0.001; drug reinstatement: interaction: F_1,17_=28.70, p<0.001, group comparisons: ext: p=0.9477, rein: p<0.001; stress reinstatement: interaction: F_1,12_=21.24, p<0.001 group comparisons: ext: p=0.9965, rein: p<0.001). AC, anterior commissure; Ext, extinction; PrL, IL, infralimbic cortex; prelimbic cortex; NAcc, nucleus accumbens core; ns, not significant; Rein, reinstatement; SA, self-administration. Data are mean ± SEM. **p<0.01, ***p<0.001.

### PrL→NAcc neurons display heterogenous activity dynamics across heroin self-administration

To assess PrL→NAcc neuronal activity dynamics across heroin self-administration and seeking, we combined our head-fixed self-administration assay with two-photon calcium imaging. We injected a retrogradely trafficked virus encoding Cre-recombinase (rgAAV2-CAG-Cre) bilaterally into the NAcc and a Cre-dependent virus encoding a calcium indicator (AAVdj-Ef1a-DIO-GCaMP6m) into the PrL, before implanting a gradient index (GRIN) lens dorsal to PrL (Fig. 2A). This allowed longitudinal imaging of individual GCaMP6m expressing PrL→NAcc neurons as mice underwent heroin self-administration (Fig 2B-D). Two-photon calcium imaging occurred during early (days 1-2), middle (days 5-6), and late (days 13-14) behavioral acquisition sessions (Fig. 2E). At the population level, PrL→NAcc neurons show an excitatory fluorescence response upon active lever pressing during early, middle and late acquisition sessions (Fig. 2F-H, top). However, when considering the calcium dynamics of individual neurons within the PrL→NAcc circuit, we observed heterogenous activity around the lever press, with some neurons excited and others inhibited (Fig. 2F-H, bottom). To determine the proportions of excited or inhibited neurons on each day, we used an area under the receiver operator characteristic (auROC) analysis that compares the average fluorescence of each neuron around the lever press (5 seconds before and after active lever press) to baseline. We observed subsets of significantly excited or significantly inhibited neurons across early, middle and late heroin self-administration (Fig. 2I, Fig. S2A-C), with significant differences in the proportion of excited neurons present on each day of heroin self-administration. Together our results suggest that distinct subsets of neurons within the PrL→NAcc circuit display opposing activity patterns during heroin self-administration.

**Figure 2.**
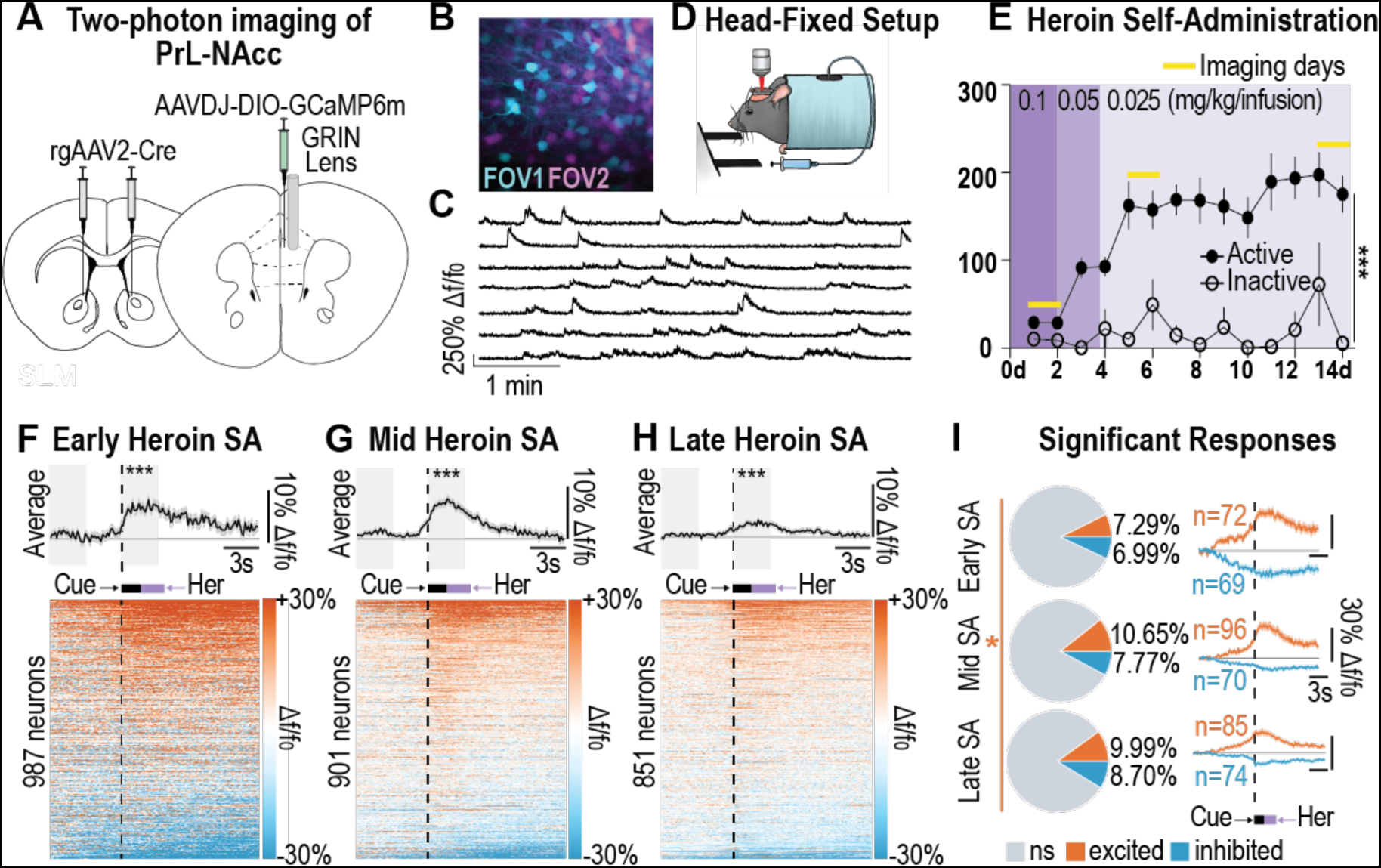
PrL→NAcc calcium activity is heterogenous across heroin self-administration. (A) Viral approach used for two-photon calcium imaging of PrL→NAcc neurons. (B) Example field of view (FOV) in cyan and second FOV in magenta separated by 50μm. (C) Example extracted signals of calcium activity of cells in FOV1 during habituation to head-fixed chamber. (D) Head-fixed apparatus used for two-photon imaging and heroin self-administration (SA). (E) Heroin SA data for imaging animals. Imaging days shown in yellow. Mice learned to press the active and not inactive lever (n=14 mice; lever: F_1,26_ = 59.45, p<0.001). (F-H) Averaged traces (top) and single-cell heatmaps (bottom) reveal PrL→NAcc activity during early (F; n=987 neurons; 14 mice), middle (G; n = 901 neurons; 14 mice), and late (H; n=851 neurons, 14 mice) SA sessions. Average fluorescence (top) increased in the 3 seconds following lever press compared to a 3 second baseline period (early: t_22_=24.94, p<0.001; mid: t_22_=31.47, p<0.001; late: t_22_=24.77, p<0.001). (I) Pie charts (left) and averaged traces (right) for each phase of heroin SA show excited (orange) and inhibited (blue) neurons with significant area under the receiver operator characteristic (auROC) scores (p<0.05). The proportion of excited neurons varied across day of heroin SA (ꭕ^2^_2_= 7.721, p=0.021) while the proportion of inhibited neurons were similar between day of heroin SA (ꭕ^2^2= 2.416, p=0.299). FOV, field of view; PrL, prelimbic cortex; NAcc, nucleus accumbens core; ns, not significant; SA, self-administration. Data are mean ± SEM. *p<0.05, **p<0.01, ***p<0.001.

### PrL→NAcc neuronal activity is heterogenous during reinstatement of heroin seeking

To measure PrL→NAcc activity dynamics during heroin seeking, following acquisition mice underwent extinction training for a minimum of 10 days where heroin and heroin-paired cues were omitted. Once active lever pressing was suppressed to less than 20% of average active presses on the last two days of heroin self-administration, mice underwent reinstatement tests in a pseudo-randomized order (Fig. 3A-C) with simultaneous two-photon calcium imaging. Mice showed elevated active lever pressing during cue- (Fig. 3A), drug- (Fig. 3B), and stress-induced (Fig. 3C) reinstatement tests compared to extinction conditions, with no changes to inactive lever presses (Fig. S3A). Average fluorescence of PrL→NAcc neurons was elevated from baseline following active lever press during cue-, drug-, and stress-induced reinstatement (Fig. 3D-F, top). However, when visualizing the fluorescence changes of individual neurons within the PrL→NAcc circuit across reinstatement tests, we observed heterogenous activity around the lever press, with some neurons excited and others inhibited (Fig. 3D-E, bottom). As described above, we then determined the proportion of significantly excited or inhibited neurons present during each reinstatement test using an auROC analysis (Fig 3G, Fig. S3B-D). We observed differences in the proportions of both excited and inhibited neurons present during each reinstatement test, with the most excited and least inhibited neurons observed during stress-induced reinstatement.

**Figure 3.**
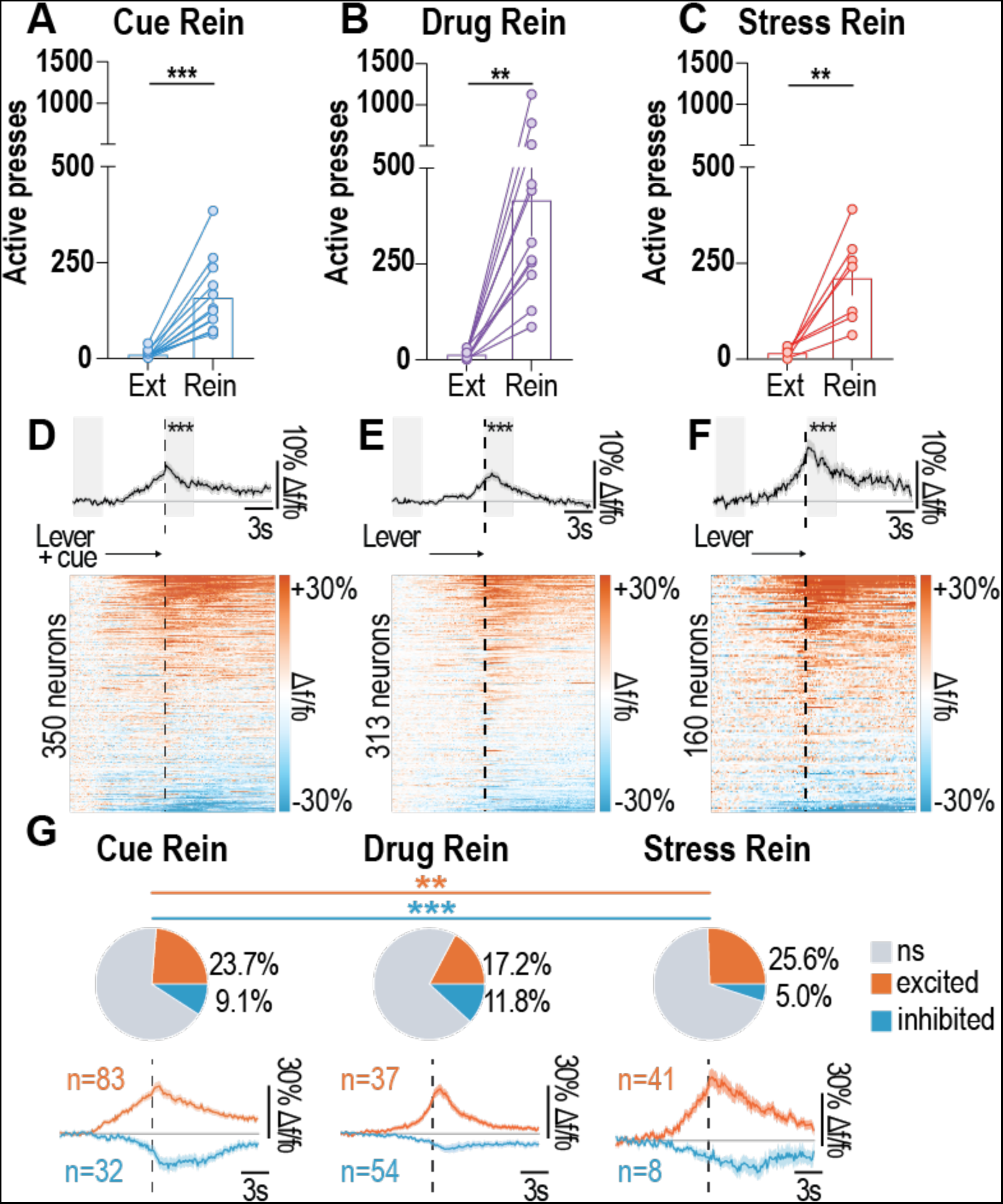
PrL→NAcc calcium activity is heterogeneous during reinstatement of heroin seeking. (**A-C**) Active lever presses during cue- (**A**), drug- (**B**), and stress-induced (**C**) reinstatement tests where active lever presses increased above previous extinction session (n=7-12 mice; cue: t11=5.867, p=0.0001; drug: t10=4.1414, p=0.0013; stress: t6=4.197, p=0.0057). (**D-E**) Averaged traces (top) and single-cell heatmaps (bottom) reveal PrL→NAcc neuronal activity during cue- (D; n=350 neurons, 12 mice), drug-(E; n=313 neurons, 11 mice) and stress-induced reinstatement (F; n=160 neurons, 7 mice). (**G-I**) Pie-charts (top) and averaged traces (bottom) for cue- (**G**), drug- (**H**), and stress-induced (**I**) reinstatement tests show excited (orange) and inhibited (blue) neurons with significant area under the receiver operator characteristic (auROC) scores (p<0.05). The proportion of excited neurons varied with reinstatement test (ꭕ^2^2=12.06, p=0.002), as did the proportion of inhibited neurons (ꭕ^2^2=19.11, p<0.001). PrL, prelimbic cortex; NAcc, nucleus accumbens core; ns, not significant; Rein, reinstatement. Data are mean ± SEM. *p<0.01, ***p<0.001.

### Spectral clustering reveals that discrete ensembles within the PrL→NAcc differentially predict aspects of heroin self-administration and heroin seeking

To identify discrete ensembles within the PrL→NAcc circuit, we combined all excited neurons (n=658) or all inhibited neurons (n=442) across each imaging session and applied a principle components analysis and spectral clustering algorithm to each population (Fig. 4A, Fig S4 A-D). This revealed 4 excited and 4 inhibited ensembles with distinct activity patterns around the active lever press (Fig. 4A). To determine whether significantly excited or inhibited neurons displayed stable activity within a behavioral session, we compared neuronal activity early in the session (first 33% of active lever presses) to activity towards the end of the session (last 33% of active lever presses) (Fig. 4B-D). For excited and inhibited neurons, we found that auROC values remained stable across early and late trials (Fig. 4C). However, when we compared early and late auROC responses for non-significant responders we observed a negative association (Fig. 4D). Together, these results suggest that significantly excited and inhibited PrL→NAcc neurons are stable classifiers of lever pressing across trials, while neurons that do not exhibit significant activity dynamics around the lever press have poor stability across trials.

**Figure 4.**
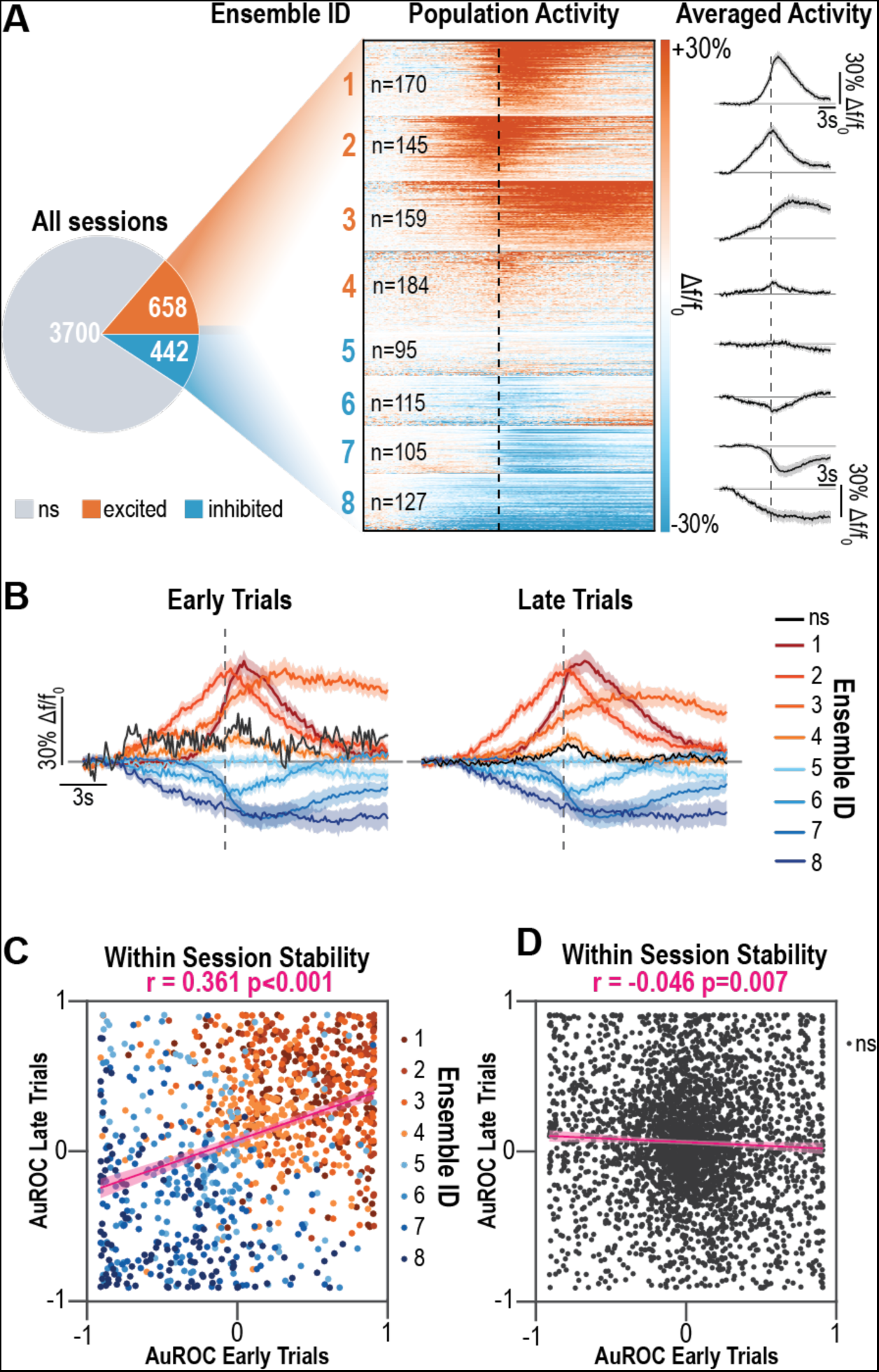
PrL→NAcc neuronal ensembles are stable across trials and differentially encode distinct aspects of heroin seeking. (**A**) Spectral clustering reveals4 excited ensembles (total n=658, orange, top half of heat map), and 4 inhibited ensembles (total n=442, blue, bottom half of heatmap) across all imaging sessions (n=8) and all animals (n=14 mice), with distinct activity patterns around the lever press (averaged line graphs, right). (B) Averaged traces for neurons that do not have significant responses (black) and for each ensemble for early trials (first 33% of trials, left) and late trials (last 33% of trials, right) (**C**) Significantly excited or inhibited neurons exhibit stable auROC scores within session (Pearson-R value=0.361, p<0.001), while (**D**) neurons that do not have significant responses exhibit negatively correlated auROC scores between early and late trials (Pearson-R value=-0.046, p=0.007). auROC, area under the receiver operator characteristic; PrL, prelimbic cortex, NAcc, nucleus accumbens core ns, not significant. Data are mean ± SEM. *p<0.05, **p<0.01, ***p<0.001

## DISCUSSION

Here we characterize the precise activity dynamics of corticostriatal projection neurons across drug self-administration and relapse. At the population level, we find PrL→NAcc projection neurons are excited during drug seeking and are necessary for relapse-like behavior. However, when we examine the activity of individual neurons, we observe heterogenous dynamics within the PrL→NAcc circuit with distinct subpopulations that display opposing activity during drug seeking. Throughout both drug self-administration and drug seeking, spectral clustering identified 4 excited and 4 inhibited PrL→NAcc neuronal ensembles, each with a distinct activity pattern around the lever press. While it remains unclear whether these ensembles discretely encode specific elements of drug seeking, such as drug-associated cues or lever pressing, our data suggest that this heterogenous activity within the PrL→NAcc circuit is required for the expression of drug-seeking behavior.

Previous investigations into the neuronal ensembles underlying drug seeking have relied on the expression of immediate early genes (IEG) to label ensembles^20–22^. Conditioned drug seeking induces the expression of the IEG *c-Fos* and its protein product Fos in a subset of prefrontal cortex neurons, and molecular silencing of these Fos-expressing ensembles confirms they play a causal role in drug seeking^20–22^. However, Fos expression requires strong and persistent neuronal activity over a period of minutes to hours^23^, and it is unclear if Fos expression is correlated with action potentials^24^. Because of this issue, studies using Fos expression to identify and target neuronal ensembles may underestimate the true number of neurons encoding a stimulus, and cannot provide information about the true computational dynamics or heterogeneity between these ensembles due to poor temporal resolution. Here we demonstrate that within the subset of PrL→NAcc neurons excited during drug seeking, there are 4 neuronal ensembles with temporally distinct activity dynamics around the lever press. These data suggest that drug-seeking ensembles identified using IEG techniques may encompass neurons with transiently different activity patterns during drug-seeking behavior. Therefore, IEG-dependent techniques likely only capture a subset of the excited ensembles we observe here, and within the labelled neurons there may be distinctly diverse activity patterns during drug seeking. These data could explain why silencing cortical IEG-labelled ensembles only modestly disrupts drug-seeking behavior^20–22^. Additionally, we find there are 4 inhibited ensembles within the PrL→NAcc circuit during drug seeking that have distinct activity patterns around the lever press. Only recently has a marker of decreased neuronal activity been identified^25^. Future studies are required to determine if activating inhibited drug-seeking ensembles in the prefrontal cortex is also sufficient to disrupt drug-seeking behavior. Corticostriatal glutamate release has long been understood to be a key feature of relapse to drug seeking across drug classes and modes of reinstatement^4–8,13,26,27^. Consistent with our results, inhibition of the PrL→NAcc circuit suppresses cue- and drug-induced cocaine seeking^5,9,10^, and aversion-resistant alcohol seeking^13^. Here, we build upon these existing studies and demonstrate that optogenetic inhibition of PrL→NAcc projection neurons prevents cue-, drug- and stress-induced reinstatement of heroin seeking, supporting the idea that PrL→NAcc circuit is a common pathway driving relapse regardless of drug class or trigger. Our single-cell calcium imaging of PrL→NAcc projection neurons reveals that different proportions of excited or inhibited neurons are recruited by each stimulus (cue, drug or stress) during drug seeking. One possibility is that each reinstatement trigger differentially recruits distinct brain regions that project to the prefrontal cortex. The PrL→NAcc neurons might then act as a critical integrator of afferent inputs, driving the expression of drug-seeking behavior in response to elevated excitatory transmission. Future research to determine whether the PrL→NAcc ensembles we have identified here can be defined by afferent inputs is warranted.

One limitation of the present study is that we have not functionally tested the contribution of each neuronal ensemble to drug-seeking behavior. Future experiments can leverage technological advances in holographic optogenetics to manipulate distinct PrL→NAcc neuronal ensembles to determine the relative contribution of each ensemble to heroin seeking^34,35^. Additionally, future studies can combine spatial transcriptomics with calcium imaging to further parse the distinct corticostriatal ensembles driving relapse. Overall, we provide the first insight into the diverse activity dynamics within the corticostriatal circuit that guide drug seeking. Further research into the anatomical and molecular features of the corticostriatal ensembles we reveal here could lead to the development of effective therapies for prevention of relapse.

## Acknowledgements

This study was funded by grants from the National Institute of Drug Abuse (NIDA): T32-DA007288 (RIG, AMW, JEP), F32-DA057794 (JEP), F32-DA053830 (EMD), K99-DA058049 (EMD), F31-DA052186 (KMV), R25-GM113278 (KTW), R01-DA051650 & R01-DA054271 (JMO), R01-DA054154 (MDS), R01-DA049711 (JFM); the National Institute on Alcohol Abuse and Alcoholism (NIAAA): T32-AA007474 (ALW), R01-AA030796 (JMO); National Center for Advancing Translational Sciences of the National Institute of Health: TL1 TR001451 & UL1 TR001450 (MRM); the Department of Veteran’s Affairs: I01BX006179 (JMO); and the MUSC College of Medicine (COMETS; JMO and MDS).

## Author Contributions

REC & JMO designed the experiments and wrote the manuscript. All authors provided technical assistance and intellectual feedback on the project.

---

**Figure S1.**
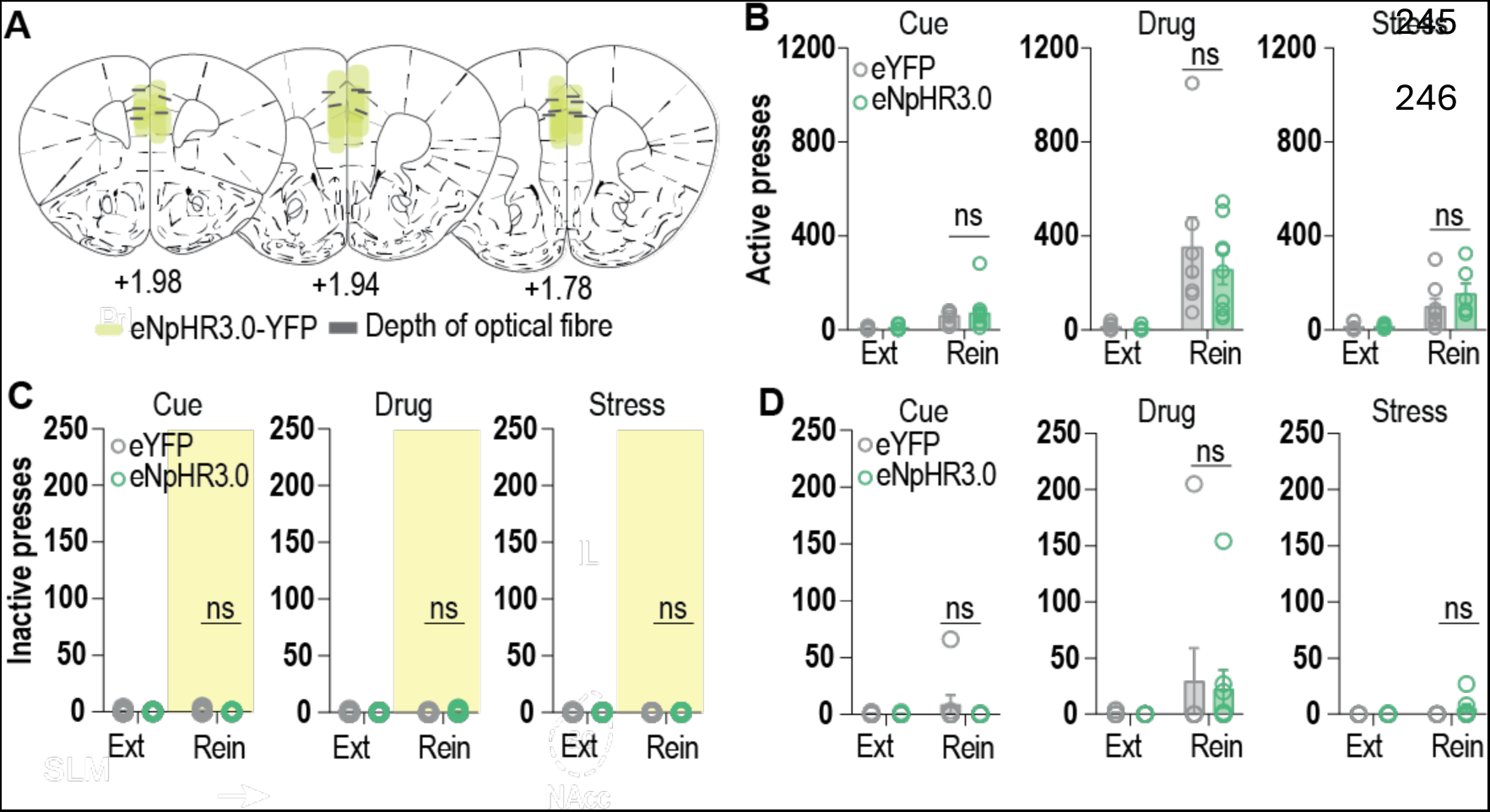
Supplemental data relating to Figure 1. Optogenetic inhibition of PrL→NAcc neurons during reinstatement of heroin seeking. (**A**) Coronal sections illustrating eNpHR3.0-YFP expression and optical fiber placements (n=9 mice). (**B**) No differences in active lever pressing under laser off conditions between eYFP and eNpHR3.0 groups during cue-, drug-, or stress-induced reinstatement tests (n=6-10/group, cue reinstatement: interaction: F_1,16_=0.067, p=0.800, group comparisons: ext: p=0.971, rein: p=0.802; drug reinstatement: interaction: F_1,14_ = 0.450, p=0.514, group comparisons: ext: p=0.996, rein: p=0.527; stress reinstatement: F_1,11_=0.957, p=0.349, group comparisons: ext: p=0.999, rein: p=0.322). (**C**) No differences in inactive lever presses under laser on conditions between eYFP and eNpHR3.0 groups during cue-, drug-, or stress-induced reinstatement tests (n=7-10/group, cue reinstatement: interaction: F_1,17_=0.607, p=0.447, group comparisons: ext: p=0.601, rein: p=0.080; drug reinstatement: interaction: F_1,17_=2.616, p=0.124, group comparisons: ext: p=0.494, rein: p=0.387; stress reinstatement: interaction: F_1,13_=1.156, p=0.302, group comparisons: ext: p=0.261, rein: p>0.999). (**D**) No differences in inactive lever presses under laser off conditions between eYFP and eNpHR3.0 groups during cue-, drug-, or stress-induced reinstatement tests (n=6-10/group, cue reinstatement: interaction: F_1,16_=1.342, p=0.264, group comparisons: ext: p=0.998, rein: p=0.204; drug reinstatement: interaction: F_1,14_=0.033, p=0.859, group comparisons: ext: p=0.999, rein: p=0.946; stress reinstatement: interaction: F_1,11_=1.706, p=0.218, group comparisons: ext: p=0.999, rein: p=0.147). PrL, prelimbic cortex, NAcc, nucleus accumbens core. Data are mean ± SEM.

**Figure S2.**
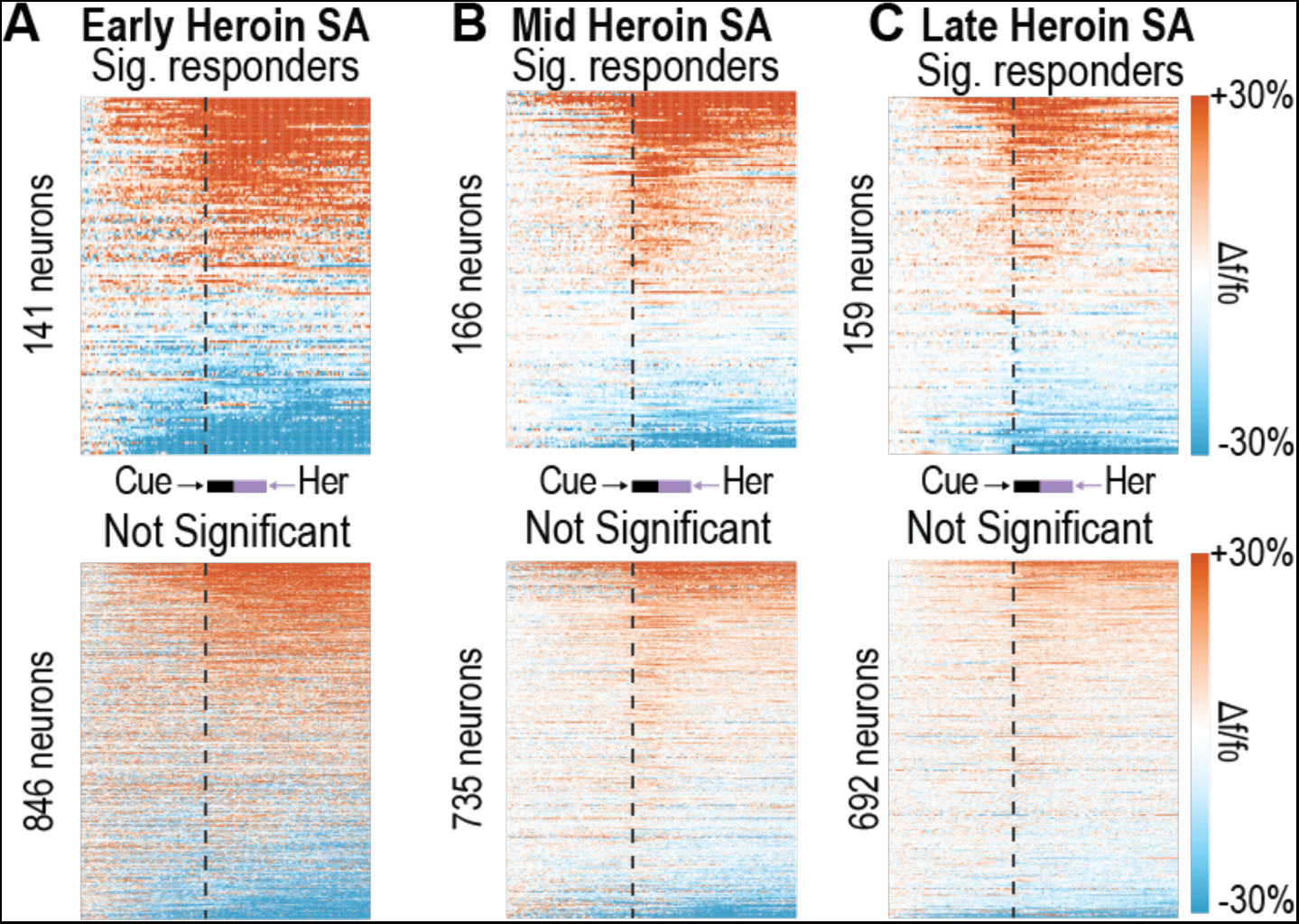
Supplemental data relating to Figure 2. Two-photon imaging of PrL→NAcc neurons during heroin self-administration. (**A-C**) Heatmaps displaying averaged fluorescent activity of each neuron across all active lever presses/session separated into significant responding neurons (top) and not significant responding neurons (bottom) for early (**A**), middle (**B**), and late (**C**) heroin self-administration (n=14 mice).

**Figure S3.**
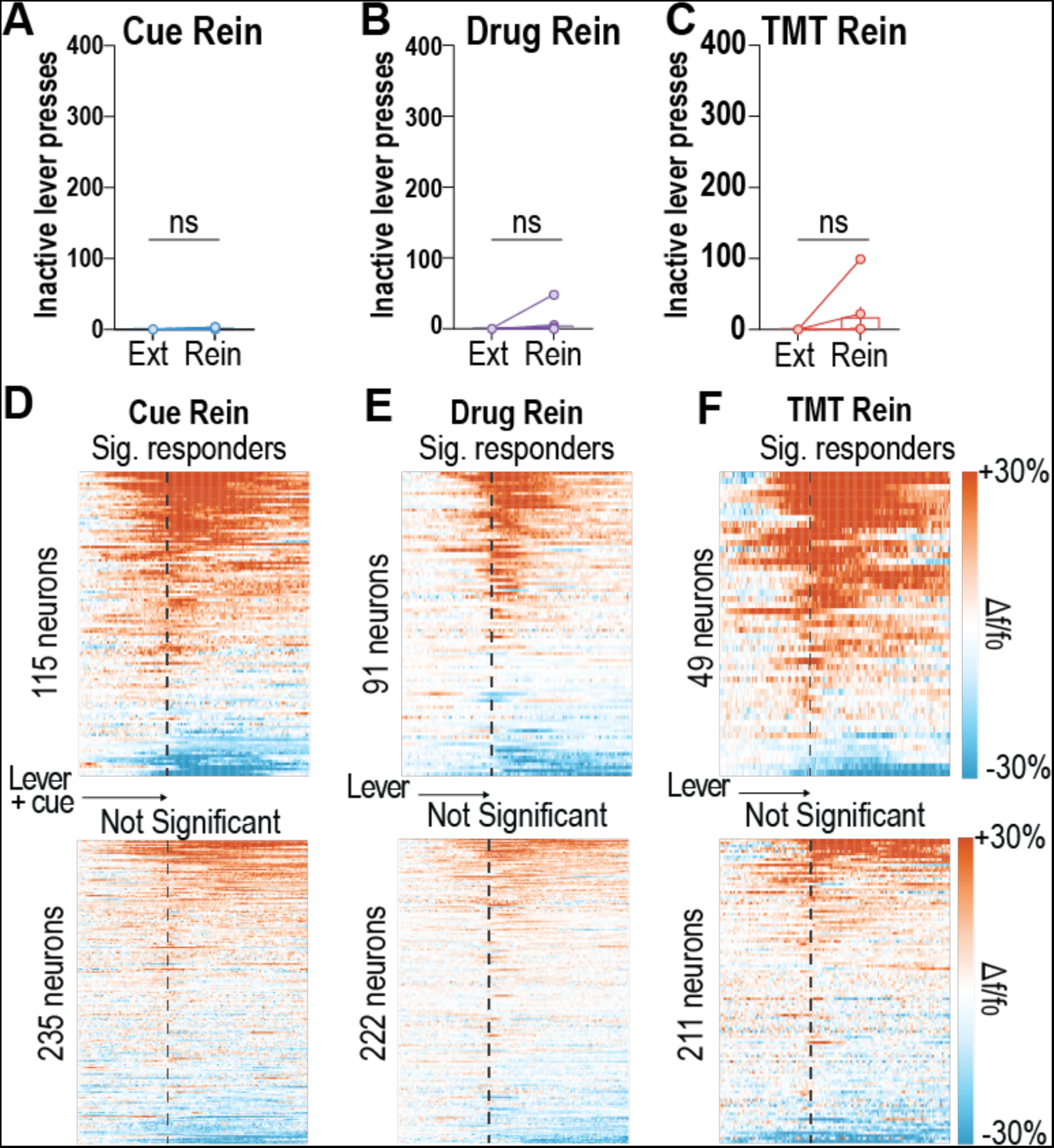
Supplemental data relating to Figure 3. Two-photon imaging of PrL→NAcc neurons during reinstatement of heroin seeking. (**A**) No differences in inactive lever presses between extinction sessions and cue-, drug-, and stress-induced reinstatement tests (n=7-12 mice, cue: t_11_=1.483, p=0.166; drug: t_10_=1.156, p=0.2746; stress: t_6_= 1.263, p=0.253). (**B-D**) Heatmaps displaying averaged fluorescent activity of each neuron across all active lever presses/session separated into significant responding neurons (top) and not significant responding neurons (bottom) for cue- (**B**), drug- (**C**), and stress-induced (**D**) reinstatement tests (n=7-12 mice). Data are mean ± SEM.

**Figure S4.**
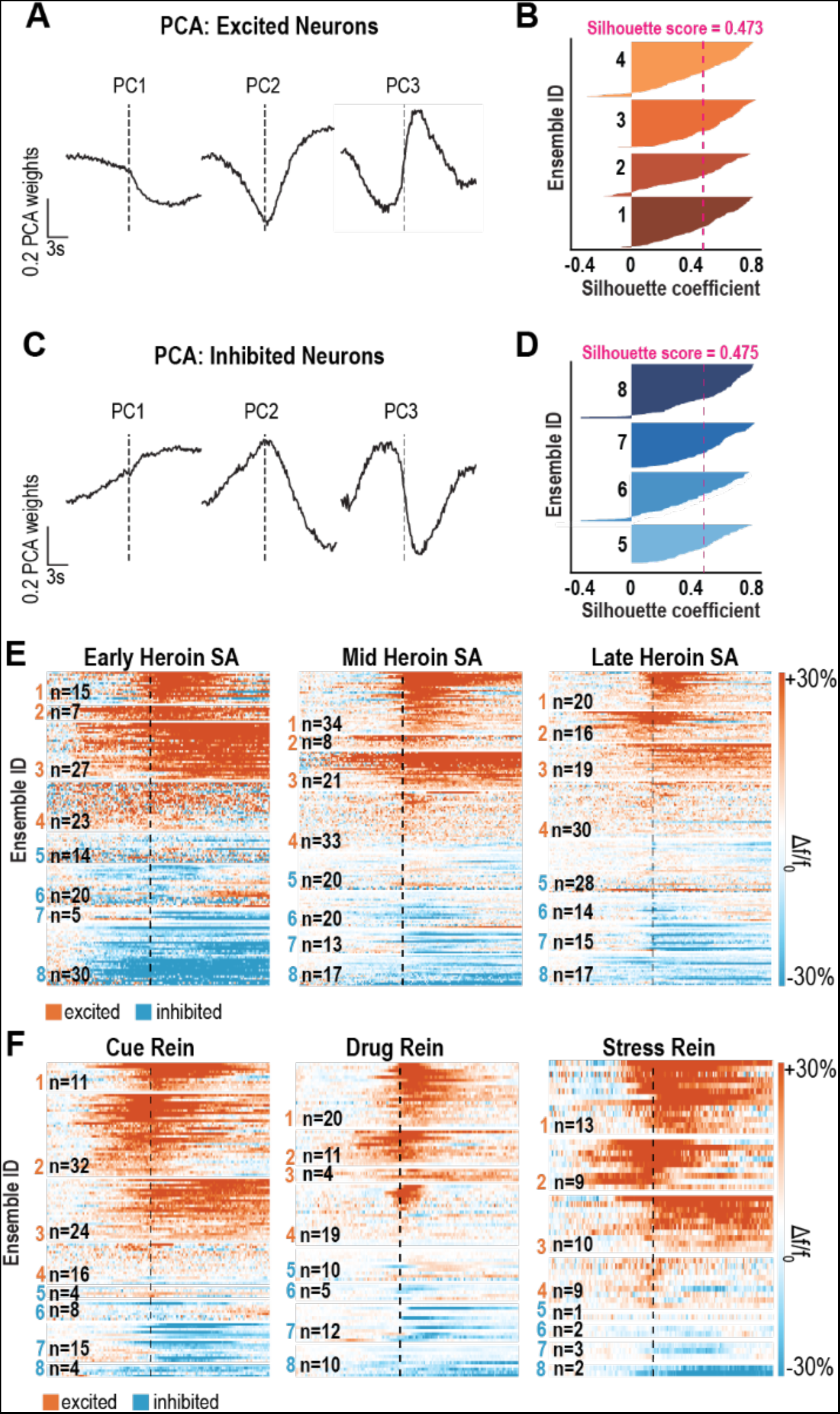
Supplemental data relating to Figure 4. Two-photon imaging of PrL→NAcc neurons and spectral clustering reveals 8 distinct neuronal ensembles present during heroin seeking. (**A-B**) Principal components analysis (**A**) and silhouette plot (**B**) show the relative fit for excited neurons from each ensemble formed by spectral clustering. (**C-D**) Principal components analysis (**C**) and silhouette plot (**D**) show the relative fit for inhibited neurons from each ensemble formed by spectral clustering. (**E-F**) Heatmaps displaying significant responding neurons split by cluter across early, mid, and late heroin self-administration (**E**) and cue-, drug-, and stress-induced reinstatement tests (**F**).

## RESOURCE AVAILABILITY

### Lead Contact

Further information and request for resources and reagents should be directed to and will be fulfilled by the lead contact, James M. Otis.

### Materials Availability

This study did not generate new unique reagents.

### Data and Code Availability

Behavioral data generated in this study, and all original code, will be deposited in the Otis Lab GitHub database and will be publicly available as of date of publication. Access to data or code prior to publication will be made upon request. Two-photon imaging datasets will be made available upon request but are not immediately available for download due to file size. Any additional information required to reanalyze the data reported in this paper is available from the lead contact upon request.

## EXPERIMENTAL MODEL AND STUDY PARTICIPANT DETAILS

### Animals

All experiments were approved by the Institutional Animal Care and Use Committee (IACUC) at the Medical University of South Carolina in accordance with the NIH-adopted Guide for the Care and Use of Laboratory Animals. Adult male and female C57BL6/J wild-type mice were group-housed pre-operatively and single-housed post-operatively, with access to standard chow and water *ad libitum* throughout all experiments. Mice were at least 8-weeks of age and 18.5g prior to study onset. Male and female mice were randomly assigned to experimental groups. Mice were housed under a reverse 12:12-hour light cycle (lights off at 8:00am), with experiments performed during the dark phase.

## METHOD DETAILS

### Surgery

For intracranial or intravenous catheter surgeries, mice were anesthetized with isoflurane (1-2.5% in oxygen; 1L/minute). Ophthalmic ointment (Akorn), topical anesthetic (2% Lidocaine; Akorn), analgesic (Ketorolac, 2 mg/kg, intraperitoneal injection), were given pre- and intra- operatively for health and pain management. An antibiotic (Cefazolin, 200 mg/kg, subcutaneous injection) was given post-operatively to reduce the possibility of infection.

#### Optogenetics surgeries

Once anesthetized, mice were placed within a stereotactic frame (Kopf Instruments). To target the PrL→NAcc circuit for optogenetic manipulations we infused a Cre-dependent virus encoding for one of two constructs (AAV5-ef1a-DIO-eNpHR3.0-eYFP; AAV5-ef1a-DIO-eYFP; 300nl/ injection site) into the PrL (AP: +1.85mm; ML: ±0.35mm; DV: −2.6mm & −2.3mm; relative to bregma), and a retrogradely-trafficked virus encoding for Cre-recombinase (rgAAV2-CAG-Cre; 400nL/side) into the anterior NAcc (AP: +1.42mm; ML: ±1.75mm; DV: −4.70mm; 10° angle). Custom-made optical fibers^30^ were implanted dorsal to the PrL injection site (AP: +1.85mm; ML: ±0.95mm; DV: −1.90mm; 10° angle), allowing laser-evoked inhibition of PrL→NAc projection neurons. A stainless-steel head ring was cemented around the optical fiber using dental cement and skull screws. Optical fiber and viral placements were confirmed post-mortem via histology (Fig. S1A).

#### Two-photon calcium imaging surgeries

Once anesthetized, mice were placed within a stereotactic frame (Kopf Instruments). To target the PrL→NAcc circuit for two-photon calcium imaging we infused a Cre-dependent virus encoding for the calcium indicator GCaMP6m (AAVdj-ef1a-DIO-GCaMP6m; 400nL/site) unilaterally into the PrL (AP: +1.85mm; ML: − 0.35mm; DV: −2.6mm & −2.3mm) and a retrogradely-trafficked virus encoding for Cre-recombinase (rgAAV2-CAG-Cre; 400nL/side) into the anterior NAcc (AP: +1.42mm; ML: ±1.75mm; DV: −4.70mm; 10° angle). A microendoscopic gradient refractive index lens (GRIN lens; 4mm long, 1mm diameter, Inscopix) was then implanted dorsal to the PrL injection site (AP: +1.85mm; ML: −0.35mm; DV: −2.25mm) allowing for chronic visualization of PrL→NAcc neurons^7,31^. Finally, a stainless-steel head ring was cemented around the GRIN lens using dental cement and skull screws. GRIN lens placement and GCaMP6m fluorescence of PrL→NAcc neurons was confirmed post-mortem.

#### IV Catheter Surgeries

Mice were allowed at least 7-days of recovery from intracranial surgery before catheterization occurred. Once anesthetized, mice were implanted with custom-made intravenous catheters, using a method previously described^9^. Catheters were implanted subcutaneously with the tubing inserted into the external jugular vein. All mice received analgesic, ophthalmic, and antibiotic treatments as described above, as well as topical antibiotic ointment and lidocaine (2%) jelly around incision cites. Following a minimum of 5-days of recovery, mice began behavioral experiments wherein catheters were flushed daily with heparinized saline (60 units/mL, 0.03mL) to maintain patency. Mice with non-patent catheters were to be excluded from the study. If necessary, patency was determined by giving mice an intravenous infusion of brevital (10 mg/mL, 0.03 mL).

### Head-fixed Behavior

#### Heroin Self-administration

Experiments involving heroin self-administration were performed as previously described^18,19^, enabling simultaneous two-photon calcium imaging^19^. After recovery from surgery, mice were habituated for 2-days to head fixation during 45-minute sessions wherein levers were not presented.

##### Acquisition

Mice next underwent heroin self-administration or saline control sessions through 14 daily sessions, during which two levers were placed in front of the animal within forelimb reach. Pressing the active lever, but not inactive lever, resulted in the presentation of a tone cue (8 kHz, 1.6s) followed immediately by the intravenous infusion of heroin (administered over a 2s epoch). A timeout period (20s) was given after each cue- and heroin-reinforced active lever press, wherein active lever pressing had no effect. Mice were trained on a fixed ratio 1 (FR1) schedule of reinforcement using a decreasing dose design (Day 1-2: 0.1 mg/kg/12.5 µL heroin, 10 infusion maximum; Day 3-4: 0.05 mg/kg/12.5 µL heroin, 20 infusion maximum; Day 5-14: 0.025 mg/kg/12.5 µL heroin, 40 infusion maximum), for a maximum of 1 mg/kg of heroin per session. To avoid issues with excessive infusion volume and overdose, mice were capped to receiving 1 mg/kg per session to prevent overdose. Self-administration sessions were a maximum of 2-hours.

##### Extinction

Following acquisition, heroin self-administering mice underwent 1-hour extinction training sessions, wherein active lever presses resulted in neither cue nor drug delivery until extinction criteria were reached. Extinction criteria were determined *a priori*, as (1) ≥ 10-days of extinction training and (2) the last 2-days of extinction training resulting in ≤20% of the average active lever pressing observed during the last 2-days of acquisition.

##### Reinstatement

After mice reached extinction criteria, they then underwent 1-hour cue-, drug-, or stress-induced reinstatement tests in a pseudorandomized order. For optogenetic experiments, each mouse experienced each reinstatement test twice, once under laser on conditions and once under laser off conditions. Between tests, mice underwent a minimum of 2 extinction sessions, until lever pressing returned to below extinction criteria for 2 consecutive days. For cue-induced reinstatement, active lever presses resulted in cue presentation as in acquisition, however drug infusions were omitted. A timeout period (20s) was given after the onset of each cue, wherein active lever pressing did not result in cue delivery. For drug-induced reinstatement, mice received an acute injection of heroin (1 mg/kg, ip) immediately before the session, and active lever presses resulted in neither cue nor drug delivery. For stress-induced reinstatement, mice were exposed to the fox feces derivative 2,5-dihydro-2,4,5-trimethylthiazoline (TMT; 30 µL; 1% v/v ddH2O) for 15-minutes, contained in a vacuum-sealed line to control duration and spread of the odorant, in the head-fixed chamber with levers removed prior to the session. TMT was then removed, and levers returned for the stress-induced reinstatement session where, like drug-induced reinstatement, active lever presses did not result in cue or drug delivery.

### Behavioral Optogenetics

We used optogenetics to inhibit the activity of bilateral PrL→NAcc neurons during cue-, drug-, or stress-induced reinstatement tests. For both eNpHR3.0 or control eYFP mice, the laser (532nm; ∼10mW) was displayed (constant light) for 30-second intervals once/minute throughout the session.

### Two-photon Calcium Imaging

We visualized GCaMP6m-expressing PrL→NAcc projection neurons using a two-photon microscope (Bruker Nano Inc) equipped with a tunable InSight DeepSee laser (Spectra Physics, laser set to 920nm, ∼100fs pulse width), resonant scanning mirrors (∼30Hz frame rate), a 20X air objective (Olympus, LCPLN20XIR, 0.45NA, 8.3mm working distance), and GaAsP photodetectors. For most animals (12/14 mice) two fields of view (FOVs) were visible through the GRIN lens (separated by >60µm in the Z-axis to avoid overlapping recordings from the same neurons), in which case we recorded from each FOV during separate imaging sessions. Data were acquired with 4 frames averaged per second using PrairieView software. Data was then converted into hdf5 format and motion corrected using SIMA^32^. Following motion correction, a motion-corrected video and averaged time-series frame were used to draw regions of interest (ROIs) around dynamic and visually distinct somas using the polygon selection tool in FIJI^33^. Fluorescent traces for each ROI were then extracted and analyzed using custom Python codes in Jupyter Notebook^7,34^. Two-photon imaging was performed during select acquisition (early: days 1-2; middle: days 5-6; late: days 13-14) and extinction sessions (early: days 1-2; late: last 2-days) and during all reinstatement tests.

### Immunohistochemistry

Free-floating 40µm coronal sections containing the PrL were blocked in 0.1M PBS with 2% Triton X-100 (PBST) with 2% normal goat serum (NGS, Jackson Immuno Research, Westgrove, PA) for 2-hours at room temperature with agitation. Sections were then incubated overnight at 4°C with agitation in GFP primary antisera diluted in 2% PBST with 2% NGS, washed 3 times for 5-minutes in PBST, then incubated in the appropriate secondary antisera diluted in PBST with 2% NGS for 4-hours at room temperature with agitation. Secondary antisera were raised in goat, conjugated to Alexa fluorophores, were used at a concentration of 1:1000, and were purchased from Invitrogen (Carlsbad, CA). Sections were then washed 3 times for 5-minutes in PBST, mounted on SuperFrost+ slides, and cover slipped with ProLong™ Gold Antifade. Slides were stored in a dark area. Brain sections were imaged using a Leica SP8 laser-scanning confocal microscope. For detection of eYFP+ cells, an OPSL 488nm laser line was used.

## QUANTIFICATION AND STATISTICAL ANALYSIS

### Behavioral Data

#### Acquisition and Extinction

Lever pressing during acquisition was analyzed across levers and days using two-way ANOVAs followed by Sidak’s post-hoc comparisons when applicable. For extinction, the first 3 days of extinction were averaged and compared to the last 3 days of extinction using a two-tailed paired t-test.

#### Reinstatement

For optogenetic experiments we compared lever pressing during the previous extinction session to lever pressing during the reinstatement test between the experimental groups (eYFP vs eNpHR3.0) using two-way ANOVAs with Sidak’s post-hoc comparisons when applicable. For two-photon imaging mice, we compared lever pressing during the previous extinction session to lever pressing during the reinstatement test using a two-tailed paired t-test. All statistical analyses were performed using GraphPad Prism statistical software. Behavioral data is represented as mean ± standard error of the mean.

### Two-photon Calcium Imaging Data

To normalize fluorescent signals, we z-scored the activity of each neuron prior to analysis. We then aligned normalized fluorescent traces of each neuron to active lever presses, including the 10-seconds beforehand, 1.6-seconds from the lever press to the start of heroin delivery, and 10-seconds after start of heroin delivery. The 21.6-second fluorescent trace was averaged across trials (active lever presses) and plotted as a peri-stimulus time heatmap across neurons. The activity of all neurons across each session were averaged and plotted as an average trace with ± standard error of the mean. The average activity of the 3-second baseline period (10-seconds prior to the lever press) was compared to the average activity 3-seconds after the lever press using a two-tailed paired t-test (scipy function: stats.ttest_rel).

To determine which individual neurons showed significant responses during the lever press period, we used an auROC analysis to compare the activity of each neuron during the 3-second baseline period to the activity during the 5-seconds before and after each lever press (with or without cue and infusion depending on session type). Significant responses with a positive auROC value were labelled as significant excited neurons and those with a negative auROC value were labelled as significant inhibited neurons. The proportions of significant excited neurons or inhibited neurons were compared across days using a chi-squared test.

We combined all significant excited neurons across all sessions and all significant inhibited neurons across all sessions into separate 2-dimensional arrays that were used to inform separate principal component analyses. The first 3 principal components were plotted into a subspace and used to inform the Scikit-learn function *sklearn.cluster.SpectralClustering*, a spectral clustering algorithm that uses a k-nearest neighbor connectivity matrix to identify unique cell clusters. Spectral clustering was chosen due to its improved performance for separating dynamic neuronal datasets as compared with other clustering algorithms^36–37^. We used spectral clustering to separately cluster excited neurons and inhibited neurons. The auROC values for each neuron were compared between early and late trials (first and last 33% respectively) using Pearson-R correlation tests with separate analyses for significant responders and not significant neurons.

